# Excess significance bias in repetitive transcranial magnetic stimulation literature for neuropsychiatric disorders

**DOI:** 10.1101/614230

**Authors:** Ali Amad, Renaud Jardri, Chloé Rousseau, Yann Larochelle, John P.A. Ioannidis, Florian Naudet

## Abstract

**Introduction:** Repetitive transcranial magnetic stimulation (rTMS) has been widely tested and promoted for use in multiple neuropsychiatric conditions, but as for many other medical devices, some gaps may exist in the literature and the evidence base for rTMS clinical efficacy remains under debate. We aimed to empirically test for an excess number of statistically significant results in the literature on rTMS therapeutic efficacy across a wide range of meta-analyses and to characterize the power of studies included in these meta-analyses.

**Methods:** Based on power calculations, we computed the expected number of “positive” datasets for a medium effect-size (standardized mean difference, SMD=0.30) and compared it with the number of observed “positive” datasets. Sensitivity analyses considered small (SMD=0.20), modest (SMD=0.50), and large (SMD=0.80) effect sizes.

**Results:** 14 meta-analyses with 228 datasets (110 for neurological disorders and 118 for psychiatric disorders) were assessed. For SMD=0.3, the number of observed “positive” studies (n=94) was larger than expected (n=35). We found evidence for an excess of significant findings overall (p<0.0001) and in 8/14 meta-analyses. Evidence for an excess of significant findings was also observed for SMD=0.5 for neurological disorders. 0 (0 %), 0 (0 %), 3 (1 %), and 53 (23 %) of the 228 datasets had power >0.80, respectively for SMDs of 0.30, 0.20, 0.50, and 0.80.

**Conclusion:** Most studies in the rTMS literature are underpowered. This results in fragmentation and waste of research efforts. The somewhat high frequency of “positive” results seems spurious and may reflect bias.

**Trial Registration:** PROSPERO 2017 CRD42017056694

## INTRODUCTION

Repetitive transcranial magnetic stimulation (rTMS) is a non-invasive neuromodulation technique, that has been increasingly used to manage drug-resistant neuropsychiatric disorders [1]. Following a seminal rTMS report in 1991[2], a myriad of potential clinical applications for neurological and psychiatric disorders quickly emerged in the literature (e.g. migraine, dysphagia, chronic neuropathic pain, depression, schizophrenia), to reach more than 3700 hits on the PubMed database in August 2018 (using “rTMS” as search term). In parallel, the publication of safety guidelines together with apparently successful proof-of-principle trials promoted the potential applications of rTMS in clinical practice. Clinics and medical centers worldwide started offering these off-label therapies [3]. As the method became more widespread, on-label treatments, particularly for depression, were progressively approved by the regulatory agencies of numerous countries, including Brazil, Israel, Australia and Canada. Based on published clinical trials, systematic reviews and meta-analyses the FDA approved the use of rTMS as a treatment for major depressive disorder (MDD), in 2008 (guidance revised in 2011); pain associated with certain migraine headaches, in 2013 [4]; and obsessive compulsive disorder in 2018 [5].

However, the evidence base for rTMS clinical efficacy remains under debate. For example, the National Institute for Health and Care Excellence (NICE) adopted more nuanced positions regarding its clinical efficacy for major depressive disorder (MDD), arguing that “the evidence on its efficacy in the short-term is adequate, although the clinical response is variable” [6] and for migraine, stating that the evidence on efficacy was limited in quantity and quality [7]. In other countries such as France, the use of rTMS in clinical practice is still not recognized by health authorities such as the Haute Autorité de Santé (HAS).

As for many other medical devices [8], some gaps may exist in the literature and rTMS may have been generally tested with less rigorous standards than drugs with the same indications. More specifically, recent empirical evaluations of the neuroscience literature suggest that reporting biases are prevalent and that most studies are underpowered [9,10]; small samples undermine the reliability of results across the field, notably due to a potential combination with reporting and publication biases. Such biases may lead to a spurious excess of statistically significant results in the literature.

In this context, we aimed to empirically test for an excess number of statistically significant results in the literature on rTMS therapeutic efficacy, across a wide range of meta-analyses and to characterize the power of studies included in these meta-analyses.

## METHODS

### Protocol and registration

We followed a protocol registered on PROSPERO (registration number: PROSPERO 2017 CRD42017056694).

### Eligibility criteria

We searched for meta-analyses gathering studies testing the efficacy of rTMS across various neuro-psychiatric conditions. We aimed to include a broad sample of meta-analyses. Meta-analyses were judged eligible when they: focused on patients with a neurological or a psychiatric condition (borderline conditions such as fibromyalgia were included and labeled as neurological); and assessed the use of rTMS regardless of the exact technical parameters employed. Only meta-analyses including randomized controlled trials (RCTs) were considered. Only comparisons with an inactive comparator (e.g. placebo or sham rTMS) were considered, regardless of the study design (parallel or cross-over). Only efficacy outcomes (clinical outcomes) were considered. If an article presented meta-analyses of different efficacy outcomes, we retained analysis of the outcome involving the largest number of study datasets.

We only retained meta-analyses in which information was provided or could be calculated per study on the number of participants in each of the two compared groups (those with the condition of interest and controls) and the standardized effect-size for the comparison (expressed as Cohen’s d, Hedges’ g, or other similar standardized metrics; binary outcomes were converted to continuous equivalent effect using Chinn transformation [11]).

Meta-analyses with less than five study datasets were excluded. This was an a priori cut off decision because it would be unlikely to make solid conclusions about the presence or absence of excess significance with limited evidence.

In case of overlapping meta-analyses on the same topic satisfying these selection criteria, the meta-analysis including the largest number of studies was retained.

### Information sources, searches and study selection process

The search was conducted on 05 February 2017 on PubMed with the following search string: “(Transcranial Magnetic Stimulation) AND Meta-analysis”. Selection was performed by two independent reviewers (AA, RJ). At a first step, references were screened based on title and abstract to identify all the relevant meta-analyses and to identify the topic of each of these publications. Then, the full text of all the remaining references was inspected to apply the selection criteria including the selection of the most comprehensive meta-analysis in each topic. In addition, and after data extraction, we compared all individual study datasets across meta-analyses to make sure that we excluded overlapping meta-analyses on the same topic. All disagreement during this selection process were resolved by consensus and consultation with a third reviewer (FN).

### Data collection process and data items

A data extraction sheet based on the Cochrane Handbook for Systematic Reviews of Interventions guidelines was developed. For each included meta-analysis, we extracted the characteristics of the meta-analysis (year, PICOS, funding), the summary measures for each meta-analysis and evidence of heterogeneity (I^2^ and Q-test). For each individual dataset included in these meta-analyses, we extracted the effect-sizes, the numbers of participants and the statistical significance of the results (i.e. p-value < .05 or not). Data collection was performed by three independent reviewers (AA, RJ, YL). All disagreement during this selection process were resolved by consensus and consultation with a fourth reviewer (FN).

### Outcome measures

Our primary outcome was the existence of an excessive significance bias among the retained meta-analyses. Our secondary outcomes corresponded to the description of the power of individual datasets; and the count of the individual meta-analyses with evidence of excess significance.

### Analysis

For each dataset in each meta-analysis, we estimated the power to detect at α = .05 an effect equal to a medium effect-size (standardized mean difference, SMD=0.30). This hypothesis was judged plausible based on the analysis of the two largest studies in MDD rated with a low risk of bias identified prior to initiating our systematic searches in a recent and comprehensive meta-analysis [12] (intention-to-treat analysis of studies by Levkovitz 2015 [13] and Leuchter 2015 [14]). Although this latter study focuses on synchronized TMS (sTMS), it was included in Brunoni et al’s meta-analysis because it was considered as very similar to rTMS in terms of clinical efficacy and acceptability. The sum of the power estimates gave the number of expected “positive” (statistically significant at p<0.05) datasets. The expected number of “positive” datasets was then compared against the observed number. We thus tested for an excess of significant findings using a binomial test in an unilateral formulation, following the method developed by Ioannidis and Trikalinos which evaluates whether there is a relative excess of significant findings possibly secondary to publication biases, selective analyses and outcome reporting, or fabricated data [15].

We performed 3 a priori defined sensitivity analyses respectively based SMD of 0.20, 0.50, and 0.80. These effect sizes were chosen a priori and based on Cohen’s classification for small, modest, and large effect sizes, respectively [16]. All quantitative data were described using medians (and min-max). The analysis was performed with R for statistical computing version 3.4.4 [17] by CR (using the libraries meta, pwr and ggplot2). Data and codes to reproduce the analyses are available on the Open Science Framework (see Supplementary data and code).

### Additional analyses

As planned in the protocol, the results were detailed separately for neurological and psychiatric disorders, and for each condition separately.

### Clarifications and amendments to the initial protocol

Before running the analysis, we had to adapt our analysis plan based on the nature of the data collected. First, there was two slightly overlapping meta-analyses on close but different topics (positive and negative symptoms in schizophrenia) with only one study in common. Since these 2 meta-analyses were not on the same topic, with different outcome measures, we included them both with the complete datasets. Second, the included meta-analyses comprised various cross-over studies. These studies were mainly treated in the meta-analyses as head-to-head comparison studies (using the first phase of treatment) and these datasets were extracted accordingly. For a minority of cross-over trials, datasets were used in the meta-analysis without considering the correlated nature of data. In this case, we checked the original results to assess the study as “positive” or not and compute the power of the study considering its cross-over design. As it was done in many included meta-analyses, for practical purposes, we considered the few 3-arm trials that we encountered as two placebo-comparisons. Last in a few cases, we identified some (3.5%) non randomised studies included in the meta-analyses. Data of these studies were kept since our aim was not to correct the initial meta-analyses.

## RESULTS

### Study selection

The searches provided a total of 215 citations. Of these, 92 studies were discarded (on the basis of title and abstract) because they did not meet the selection criteria. After examination of the full text of the remaining 123 articles, 109 additional references were discarded. 14 meta-analyses (8 for neurological disorders[18–25] and 6 for psychiatric disorders[12,26–30]) were included in the analysis. A flowchart, detailing the study selection process and reasons for exclusion, is provided in **Figure 1**.

**Figure 1:**
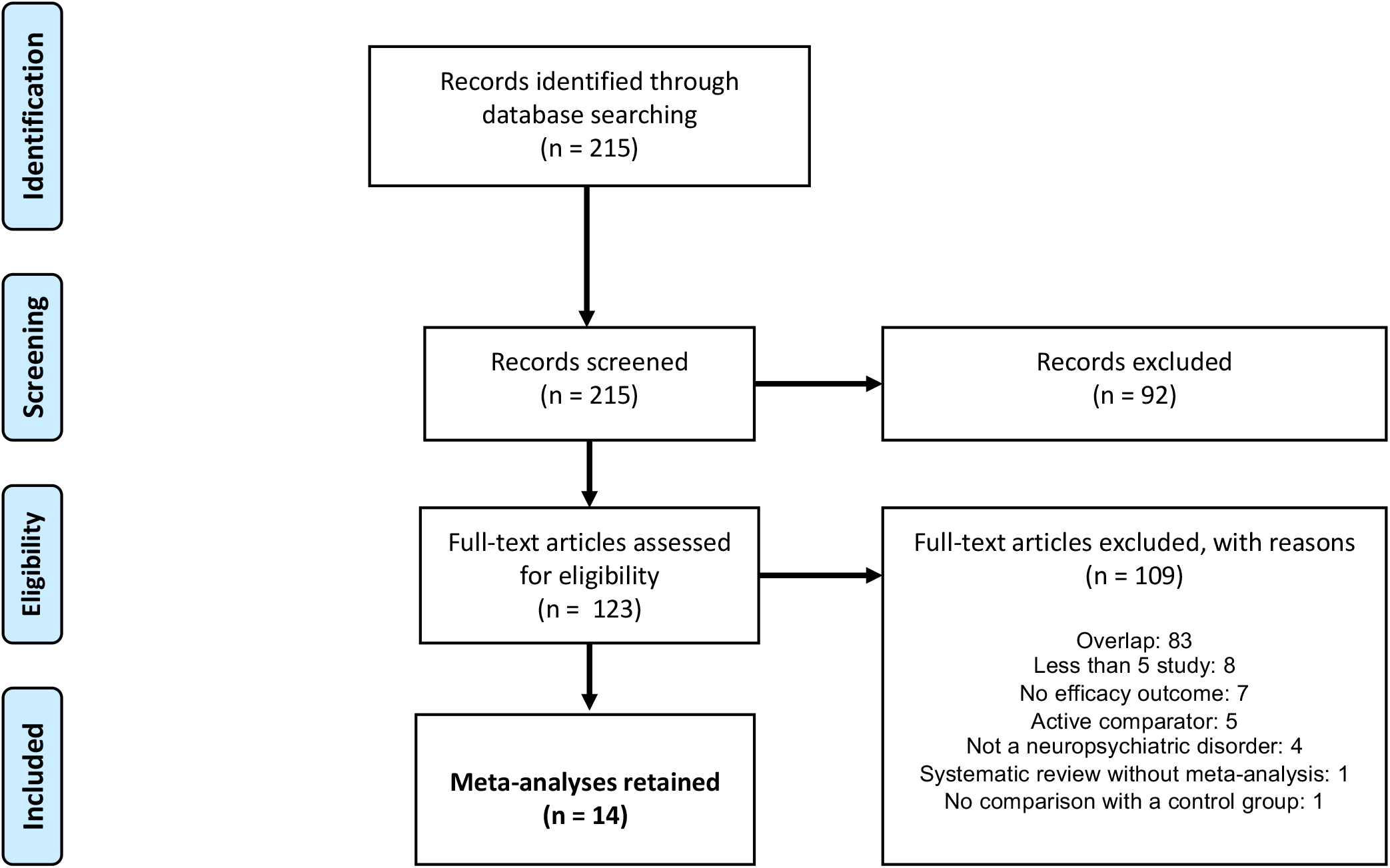
Flow chart of literature search.

### Meta-analyses characteristics

The 14 included meta-analyses represent a total of 228 datasets (110 for neurological disorders and 118 for psychiatric disorders). Of those 8 non-randomised studies were identified (7 for neurological disorders and 1 for psychiatric disorders). The median number of datasets per meta-analysis was 11.5 (min-max: 5-50), respectively 10.5 (min-max: 5-25) for neurological disorders and 13.5 (min-max: 8-50) for psychiatric disorders). All individual datasets are available on the Open Science Framework: 183 were parallel datasets (79 for neurological disorders and 104 for psychiatric disorders) with a median sample size of 27 (min-max: 10-301) and 45 were cross-over datasets (31 for neurological disorders and 14 for psychiatric disorders) with a median sample size of 12 (min-max: 4-46).

In 13/14 meta-analyses, there were nominally statistically significant differences between the active and control groups (7/8 and 6/6 in neurological and psychiatric conditions respectively). 11 effects-sizes had an absolute magnitude (SMD) exceeding 0.50. There was nominally statistically significant heterogeneity (p<0.10) in 9/14 meta-analyses (not mentioned in one). I^2^ values exceeding 50% were noted in 7/14 meta-analyses, and 2 of those had values exceeding 75%. **Table 1** details characteristics of each included meta-analysis.

**Table 1:** Description of included meta-analyses.

| Meta-analysis | Topic | Number of parallel/cross-over data sets | Median (range) sample size in parallel studies* | Median (range) sample size in cross-over studies | Summary standardized mean different [95% CIs] | I<sup>2</sup>, Qtest p-value < 0.10 | Outcomes |
| --- | --- | --- | --- | --- | --- | --- | --- |
| **Neurology** |  | **79/31** | **27 (10-150)** | **12 (4-46)** |  |  |  |
| Ren CL et al. 2014 | Aphasia in Stroke Patients | 5/0 | 21 (10-29) |  | **1.26 [0.80; 1.71]** | 0 / NO | Severity of aphasia impairment<br>Or Expressive language<br>Or Receptive language |
| Liao X et al. 2015 | Cognitive Impairment in Alzheimer's disease | 4/6 | 18 (10-22) | 9 (7-12) | **1.00 [0.41; 1.58]** | 0.68 / YES | Cognition |
| Jin Y et al. 2015 | Chronic neuropathic pain | 8/17 | 28.5 (11-70) | 14 (11-46) | **0.86 [0.56; 1.15]** | 0.81 / NA | Pain level |
| Liao X et al. 2016 | Dysphagia after stroke | 9/0 | 22 (18-29) |  | **1.24 [0.67; 1.81]** | 0.65 / YES | Swallowing function and dysphagia |
| Chung CL et al. 2016 | Motor signs in Parkinson's disease | 10/8 | 24 (13-98) | 10 (4-21) | **0.31 [0.11; 0.51]** | 0.32 / YES | Physical function and motor signs |
| Hou WH et al. 2016 | Fibromyalgia | 11/0 | 26 (10-54) |  | **0.70 [0.47; 0.94]** | 0.24 / NO | Pain |
| Graef P et al. 2016 | Function after stroke | 8/0 | 18.5 (11-66) |  | 0.03 [-0.25; 0.32] | 0 / NO | Upper-limb motor function recovery |
| Shen X et al. 2017 | Post- stroke depression | 24/0 | 75 (32-150) |  | **1.25 [0.96 ;1.54]†** | 0.96 / YES | Depression severity |
| **Psychiatry** |  | **104/14** | **26.5 (10-301)** | **14.5 (9-32)** |  |  |  |
| Slotema CW et al. 2013 | Auditory verbal hallucinations | 15/10 | 26 (11-50) | 15.5 (10-18) | **0.44 [NA] p < 0.001** | 0.27 / YES | Auditory verbal hallucinations,<br>Or Psychosis severity |
| Shi C et al. 2014 | Negative symptoms in schizophrenia | 12/0 | 21 (10-40) |  | **0.53 [0.19; 0.87]** | 0.51 / YES | Negative symptoms severity |
| Trevizol AP et al. 2016 | Post traumatic stress disorder | 7/1 | 18 (14-20) | 9 (9-9) | **0.74 [0.06; 1.42]** | 0.71 / YES | Severity |
| Trevizol AP et al. 2016 | Obsessive compulsive disorder | 15/0 | 23 (18-65) |  | **0.43 [0.74; 0.13]††** | 0.58 / YES | Obsessive compulsive symptomatology |
| Maiti R et al. 2016 | Craving in substance use disorder | 7/1 | 29 (18-63) | 14 | **0.75 [0.29; 1.21]** | 0.34 / NO | Craving intensity |
| Brunoni AR et al. 2017 | Major depressive disorder | 48/2 | 32.5 (15-301) | 22 (12-32) | **0.59 [0.39; 0.78]†††** | 0.48 / YES | Remission |

\* Cross over studies analyzed as parallel studies were considered as parallel studies
† The initial meta-analysis reported mean differences. This SMD was computed using a random effect model using the values reported in the paper.
†† The initial meta-analysis reported mean differences (wrongly presented as SMD). This SMD was computed using a random effect model using the values found in previous meta-analyses and/or original studies since the initial meta-analysis did not report enough details and the data were not available after contacting the study author.
††† Study data sets were presented as odds ratio. These were converted to SMD and pooled using a random effect model to compute this effect size.

### Observed versus expected number of “positive” study datasets (main analysis)

94 “positive” study datasets were observed across the 218 datasets included with a higher proportion for neurological than psychiatric disorders (61/110 versus 33/118). Under the main assumption (SMD=0.3), in all 14 meta-analyses, the number of observed “positive” studies (n=94) is larger than expected (n=35). We found evidence for an excess of significant findings overall (p<0.0001) and specifically in 8/14 meta-analyses.

### Observed versus expected number of “positive” study datasets (sensitivity analysis)

Results were similar in the sensitivity analysis based on the assumption of a small effect-size (SMD=0.2), where only 22 significant “positive” datasets would have been expected instead of the 94 observed (p<0.0001). Evidence for an excess of significant findings was also observed (p=0.0028) under the assumption of a medium effect size (SMD=0.5). In 8/14 meta-analyses, the number of observed positive studies is larger than expected (**Table 2**). Conversely, under the assumption of a large effect size (SMD=0.8), the expected “positive” datasets (n=136) would suggest no excess significance either overall or for any single meta-analysis.

**Table 2:** Excess significance testing, overall, by specialty field (neurological disorders, psychiatric disorders) and across all meta-analyses.

| Meta-analysis | Topic | Number of individual studies | Observed number of positive studies | Expected number of positive results (SMD = 0.3) | p-value (SMD =0.3) | Expected number of positive results (SMD = 0.2) | p-value (SMD = 0.2) | Expected number of positive results (SMD = 0.5) | p-value (SMD = 0.5) | Expected number of positive results (SMD = 0.8) | p-value (SMD=0.8) |
| --- | --- | --- | --- | --- | --- | --- | --- | --- | --- | --- | --- |
| Ren CL et al. 2014 | Aphasia in stroke patients | 5 | 3 | 0.48 | 0.0074 | 0.35 | 0.0031 | 0.89 | 0.0425 | 1.86 | 0.2694 |
| Liao X et al. 2015 | Cognitive impairment in Alzheimer's disease | 10 | 6 | 1.14 | 0.0003 | 0.78 | < 0.0001 | 2.31 | 0.0133 | 4.81 | 0.3308 |
| Jin Y et al. 2015 | Chronic neuropathic Pain | 25 | 18 | 5.19 | < 0.0001 | 2.99 | < 0.0001 | 11.07 | 0.0047 | 18.92 | NA |
| Liao X et al. 2016 | Dysphagia after stroke | 9 | 6 | 0.95 | 0.0001 | 0.67 | < 0.0001 | 1.89 | 0.0039 | 4.05 | 0.1651 |
| Chung CL et al. 2016 | Motor signs in Parkinson's disease | 18 | 3 | 2.50 | 0.4668 | 1.60 | 0.2119 | 5.27 | NA | 10.15 | NA |
| Hou WH et al. 2016 | Fibromyalgia | 11 | 3 | 1.19 | 0.1069 | 0.83 | 0.0446 | 2.35 | 0.4265 | 4.84 | NA |
| Graef P et al. 2016 | Motor function after stroke | 8 | 0 | 0.87 | NA | 0.60 | NA | 1.71 | NA | 3.40 | NA |
| Shen X et al. 2017 | Post- stroke depression | 24 | 22 | 5.87 | < 0.0001 | 3.25 | < 0.0001 | 13.04 | < 0.0001 | 21.00 | 0.4075 |
| **All neurological disorders** |  | **110** | **61** | **18.18** | **< 0.0001** | **11.07** | **< 0.0001** | **38.52** | **< 0.0001** | **69.02** | **NA** |
| Slotema CW et al. 2013 | Auditory verbal hallucinations | 25 | 5 | 3.64 | 0.2955 | 2.30 | 0.0733 | 7.88 | NA | 15.54 | NA |
| Shi C et al. 2014 | Negative symptoms in schizophrenia | 12 | 4 | 1.26 | 0.0305 | 0.90 | 0.0093 | 2.49 | 0.2253 | 5.24 | NA |
| Trevizol AP et al. 2016 | Post traumatic stress disorder | 8 | 3 | 0.76 | 0.0337 | 0.56 | 0.0147 | 1.43 | 0.1591 | 3.019 | NA |
| Trevizol AP et al. 2016 | Obsessive compulsive disorder | 15 | 3 | 1.93 | 0.3032 | 1.27 | 0.1275 | 4.05 | NA | 8.18 | NA |
| Maiti R et al. 2016 | Craving in substance use disorder | 8 | 1 | 1.08 | NA | 0.7 | 0.5181 | 2.30 | NA | 4.61 | NA |
| Brunoni AR et al. 2017 | Major depressive disorder | 50 | 17 | 8.36 | 0.0023 | 5.14 | < 0.0001 | 16.95 | 0.5469 | 30.94 | NA |
| **All psychiatric disorders** |  | **118** | **33** | **17.04** | **0.0001** | **10.85** | **< 0.0001** | **35.11** | **NA** | **67.52** | **NA** |
| **Global analysis** |  | **228** | **94** | **35.22** | **< 0.0001** | **21.92** | **< 0.0001** | **73.63** | **0.0028** | **136.54** | **NA** |

NA: test not applicable: the expected number of “positive” studies is larger than the observed number of “positive” studies

### Power of individual datasets

The distributions of all computed dataset powers under the 4 different assumptions for effect-size are presented in **Figure 2**. 0 (0 %), 0 (0 %), 3 (1 %), and 53 (23 %) out of the 228 individual datasets had a calculated power > 0.80, respectively for SMDs of 0.30, 0.20, 0.50, and 0.80.

**Figure 2:**
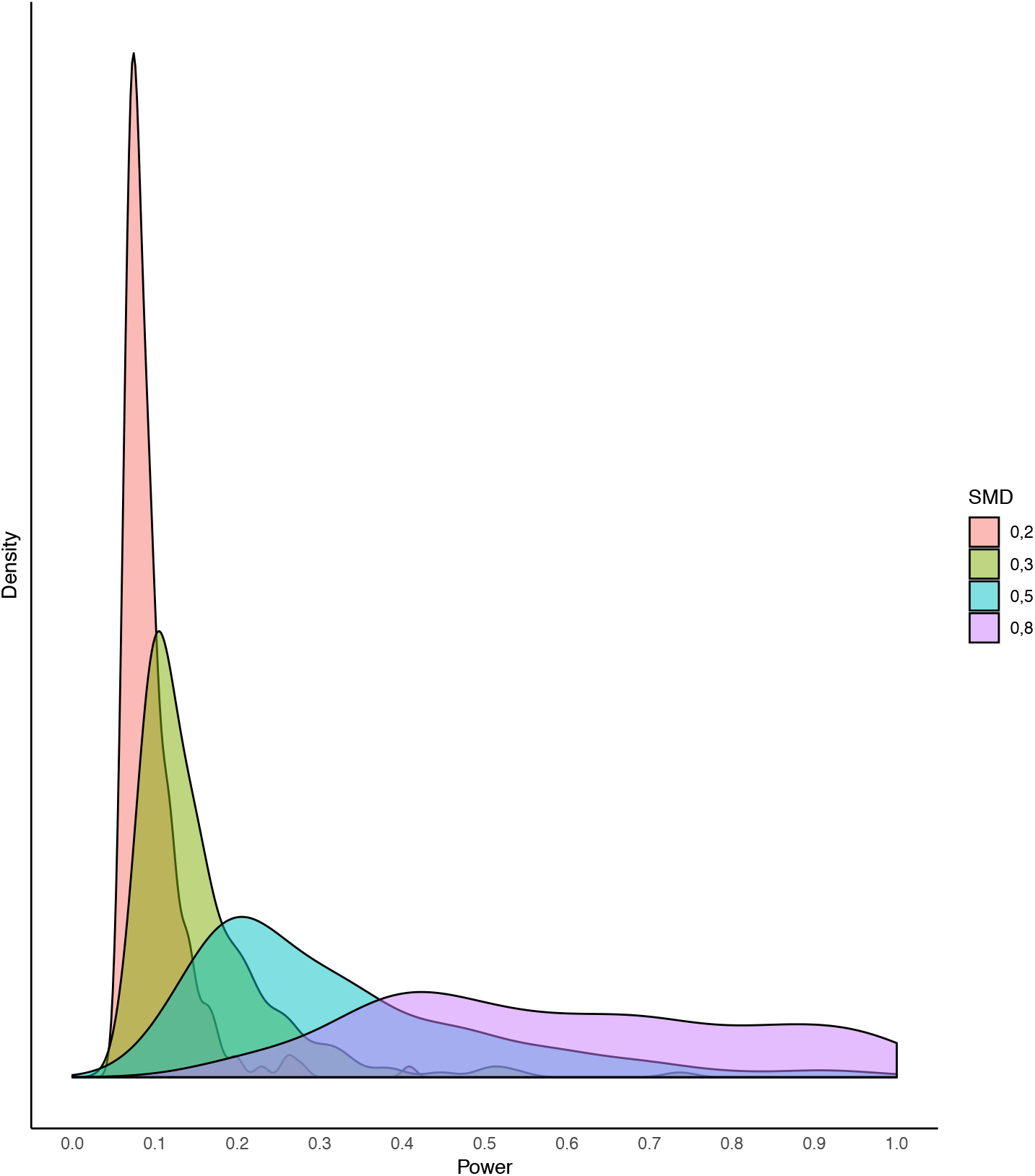
Distribution plots of datasets power under different hypothesis of rTMS true effect.

### Additional analysis

For meta-analyses of neurological disorders, evidence for an excess of significance was found for a SMD of 0.2 (Observed = 61; Expected = 11; p-value < 0.0001), 0.3 (Observed = 61; Expected = 18; p-value < 0.0001), or 0.5 (Observed = 61; Expected = 38; p-value < 0.0001), but not 0.8.

For meta-analyses of psychiatric disorders, evidence for an excess of significance was found for a SMD of 0.2 (Observed= 33; Expected = 11; p-value < 0.0001), or 0.3 (Observed = 33; Expected = 17; p-value = 0.0001), but not 0.5 or 0.8. **Table 2** details these results between meta-analyses of neurological and psychiatric disorders.

## DISCUSSION

Based on our literature search, we identified 14 meta-analyses comparing rTMS versus an inactive comparator in various neurological or psychiatric disorders. All these published meta-analyses except one (motor function after stroke) reported evidence that rTMS was an effective treatment in aphasia in stroke patients, cognitive impairment in Alzheimer’s disease, chronic neuropathic pain, dysphagia after stroke, motor signs in Parkinson’s disease, fibromyalgia, post-stroke depression, auditory verbal hallucinations, negative symptoms in schizophrenia, post-traumatic stress disorder, obsessive compulsive disorder, craving in substance use disorder and MDD. A re-evaluation of the 218 datasets taken from these 14 meta-analyses found 94 (43%) “positive” datasets (61/110 for neurological and 33/118 for psychiatric disorders). However, our analyses suggest that this number is too large if, overall, the “true” effect of rTMS was small (e.g., 21 would be expected for a small SMD of 0.2) or medium (e.g., 35 positive datasets would be expected for an SMD of 0.3 and 73 would be expected for a SMD of 0.5). This excess of significant results may be mostly driven by meta-analyses conducted in neurological disorders, while no excess of significance was detected for psychiatric disorders under the assumption of a SMD of 0.5.

Excess significance has been described for various therapeutic interventions including interventions for neurological and mental disorders, such as antidepressants [31] or psychotherapies [32]. Here, we report suggestive evidence of this phenomenon also in the rTMS literature for these conditions. Evidence for an excess of significance was not robust in our last sensitivity analysis, which assumed that the rTMS effect-size was indeed large. However, one would have to be very optimistic about the general merits of this intervention to assume such large benefits. It is more likely that combination of bias (poor research design and poor data analysis) and selective outcome reporting generally encourages false-positive findings and often disturbs the balance of findings in favor of “positive” ones [33] and can give the impression of some very large benefits.

Importantly, we found that the average statistical power of individual studies in the rTMS literature is very low. Indeed, while a power of 80% or higher is often considered as conventional in RCTs [34,35] we only found 23 % of the included datasets with a power > 80% to detect a large (and rather implausible) effect-size of rTMS. Only 3 datasets (1%) had sufficient power to detect a more plausible, modest effect size (0.5).

These findings are consistent with the average statistical power of studies in neurosciences that has been previously described as being very low [9,10,36]. Well described consequences of this include overestimates of effect sizes [9] and low reproducibility of results as suggested also by our main analysis. In addition, and despite the “non-invasive” character of the rTMS method, ethical issues arise when it comes to testing such interventions in human subjects. In other words, including participants in underpowered studies is not solely wasteful, but appears to unnecessarily expose participants to adverse events (e.g. headaches). Even if adverse events are not common, wasted time and effort to participate in research protocols would not be justified. Eventually, when these results are translated in clinical practice, excess of significance may result in a wrong evaluation of benefit-risk ratio for individual patients. This may raise questions for regulators, as illustrated by the divergent positions between FDA, NICE and HAS. It can also lead to potentially disproportionate hope in patients and dilemmas for clinicians who may want to use these strategies. Finally, the care that is offered to patients is liable to be questioned and/or discredited.

Excess of significance testing is exploratory by nature [15]. It must not be interpreted as providing a firm answer, but it rather suggests the existence of potential biases. The test depends critically on the assumption one has chosen considering the “true” effect-size. Previous studies [9,36] have often considered that this “true” effect-size might be approximated by each meta-analysis’ summary estimates, but this is affected by potential biases and thus it may be exaggerated. Therefore, the largest study is often used as an indicator of the true effect-size since it is considered to be more unbiased. However, this requires the existence of some large enough study. In the case of rTMS, when we were planning the study, preliminary looks at various meta-analyses suggested that the concept of “largest study” was not applicable in these meta-analyses that ubiquitously included only small or very small studies. The largest study would not be large enough to put much more trust on it than the others. 12/14 meta-analyses involved no dataset with more than 100 participants (including 6 meta-analyses involving no dataset with more than 50 participants).

We therefore used as a common reference a plausible effect-size derived from the two largest studies without risk of bias that we identified in MDD. We choose this topic (MDD) because it was the most extensively studied and accepted for rTMS efficacy: rTMS has an official approval from the FDA [37] and, even if more nuanced, the NICE acknowledged that there is a benefit on the short-term [6]. In addition, we performed a series of planned sensitivity analysis to explore the robustness of our initial finding. To make our judgement on the importance of each effect-size, we relied on Cohen’s classification defining small, modest and large effect-sizes [16].

As mentioned above, some meta-analyses found very large effect sizes for rTMS with 4 meta-analyses showing point estimates > 1. Most likely excess significance driven by small studies has produced these large effect sizes that are well above the usual effect sizes generally observed in the medical literature, usually around 0.40 and rarely exceeding 1 [38]. An empirical analysis of 136,212 clinical trials between 1975 and 2014 extracted from meta-analyses from the Cochrane database of systematic reviews found a median Cohen’s d of 0.20 (0.11–0.40) and effect sizes of 0.8 were rare [39]. In fact, very large treatments effect in small studies rarely appear to be a reliable marker for a benefit that is reproducible and directly actionable [40].

Our study provides a bird’s eye view on this literature without considering subtleties in terms of stimulated zone and stimulation parameters, as it was performed in the meta-analyses included in our analyses. Such differences, in theory, may generate genuine heterogeneity in effect sizes and genuine heterogeneity may also exist for different clinical settings, populations, and indications. An obvious next step would be to describe all individual studies separately with both all stimulation parameters and risks of bias. Understanding genuine heterogeneity across patients and clinical settings would nevertheless require much larger studies than those conducted to-date.

Because we aimed to be comprehensive and to avoid duplication of individual studies, we only retained the largest meta-analysis in each topic. This was necessary given the number of overlapping meta-analyses on the same topic we found. However, we may have missed some studies only included in smaller meta-analyses. In addition, due to the unavoidable delay between literature searches and publication of meta-analyses and between our own literature search and publication of this study, we may have missed some recent, modest size studies such as for example a recent (and “negative”) study of 164 patients in MDD [41].

Most (or even all) studies in the rTMS literature are underpowered. This results in fragmentation and waste of research efforts. The number of “positive” trials is substantially overestimated if the true effect size of rTMS is small to modest. We call for large and collaborative studies in the field that would help dissect whether bias is responsible for most if not all of the benefits observed, or there are still important benefits that can be reaped from rTMS in specific circumstances. The current appearance of the evidence as being strongly favorable for almost every condition where this is intervention has been tried is too good to be true.

## STATEMENTS

### Author Contributions

FN, JPAI, AA and RJ conceived and designed the experiments. AA, RJ, YL, FN performed the experiments. CR analyzed the data. CR and FN interpreted the results. FN and AA wrote the first draft of the manuscript. RJ, CR, YL, JPAI contributed to the writing of the manuscript. AA, RJ, CR, YL, JPAI and FN agreed with the results and conclusions of the manuscript. All authors have read, and confirm that they meet, ICMJE criteria for authorship. All authors had full access to all of the data (including statistical reports and tables) in the study and can take responsibility for the integrity of the data and the accuracy of the data analysis. AA is the guarantor.

